# Natural field diagnosis and molecular confirmation of fungal and bacterial watermelon pathogens in Bangladesh: A case study from Natore and Sylhet district

**DOI:** 10.1101/2024.07.03.601980

**Authors:** Raihan Ferdous

**Author notes:** Corresponding author: Raihan Ferdous.

## Abstract

The study investigated watermelon diseases in Sylhet and Natore Districts of Bangladesh, characterized by contrasting climatic conditions. Sylhet experiences lower temperatures and high rainfall, while Natore has higher temperatures and low rainfall. In these survey regions, 40 watermelon fields were selected, and 10 diseases were observed, including 4 fungal, 3 bacterial, 2 water mold, and 1 viral disease. The observed diseases were Anthracnose, Cercospora leaf spot, Fusarium wilt, Gummy stem blight, Downy mildew, Phytophthora fruit rot, Bacterial fruit blotch, Angular leaf spot, Yellow vine, and Watermelon mosaic disease. Molecular analysis was done in the Plant Pathology Lab at Sher-e-Bangla Agricultural University using the specific primers for fungal (ITS1/ITS4) and bacterial (27F/1492R) DNA regions and identified nine pathogen species, excluding the causal organism of the viral disease. The identified pathogens included *Colletrotrichum orbiculare, Cercospora citrullina, Fusarium oxysporum, Stagonosporopsis cucurbitacearum, Pseudoperonospora cubensis, Phytophthora capsici, Acidovorax citrulli, Pseudomonas syringae,* and *Serratia marcescens*. The sequencing of the identified pathogens revealed high homology (98.91-99.71%) with known sequences in the GenBank database. Phylogenetic analysis showed six clusters for fungal and water mold pathogen isolates and three for bacterial isolates where the percentages of replicate trees were 100% in all the cases. Among the identified diseases, the highest disease occurrence showed by Fusarium wilt (47.5%) following Gummy stem blight (41.5%) in the Sylhet region while Angular leaf spot (37.5%) followed Yellow vine (33%) in the Natore area. Fusarium wilt also showed high disease intensity showcasing its devastating impact on yield. The study highlights the influence of environmental conditions on disease prevalence and underscores the need for tailored management strategies. These findings provide a foundation for developing targeted disease management practices for sustainable watermelon cultivation in Bangladesh.

## Introduction

Bangladeshi people love to eat watermelon (*Citrullus lanatus*) in the summer season. This significant horticulture crop is cultivated in warm locations across the globe and is prized for its juicy, sweet fruit [1]. In Bangladesh, watermelon is a major summer cash crop with high domestic demand. Farmers receive the majority of their income from the marketing and distribution of watermelon [2]. Bangladesh is seeing a daily rise in the demand for watermelon consumption. Regarding earlier decades, the state of production also improved. At present, the production of this fruit was 550,000 and 556,000 metric tons in 2021-2022 and 2022-2023 respectively [3]. It is currently produced commercially in Bangladesh, where growers can trade it for a substantial amount of foreign exchange. Therefore, watermelon production can be vital to the nation’s growth in the economy [4].

Like other crops, controlling diseases and pests are major factors in watermelon production. In particular, diseases have a significant impact on crop quality and yield. Several fungal, bacterial, viral, and nemic diseases can affect watermelon. These include Angular leaf spot (*Pseudomonas syringae*), Bacterial fruit blotch (*Acidovorax citrulli*), Bacterial leaf spot (*Xanthomonas campestris*), Bacterial soft rot (*Erwinia carotovora*), Alternaria leaf spot/blight (*Alternaria cucumerin*), Anthracnose (*Colletotrichum orbiculare*), Phytophthora fruit rot (*Phytophthora capsici*), Downy mildew (*Pseudoperonospora cubensis*), Fusarium wilt (*Fusarium oxysporum*), Fusarium fruit rot (*Fusarium equiseti*), Gummy stem blight (*Stagonosporopsis cucurbitacearum*), and so on [5–6]. When crops get infected, Fusarium wilt and Phytophthora fruit rot are the most destructive diseases to watermelon. Furthermore, foliar diseases that affect the crop annually, like Powdery mildew, Gummy stem blight, and Anthracnose, force growers to invest extensively in crop protection and agricultural practices for managing these diseases [7].

At Gurudashpur and Baraigram in Natore, farmers cultivate watermelon as an intercrop with garlic [8]. It grows widely throughout the region from April to June, and with the right care, it can be grown all year round [9]. On the other hand, at Jaintiapur and Gowainghat in Sylhet district, farmers produce watermelon as an early variety to protect their crops from flooding. They get comparatively high values as an off-season product. After getting significant benefits farmers are now becoming more interested in watermelon farming in their region [10]. These two districts of Bangladesh are now increasingly contributing to watermelon production and uncovering the promise of exporting watermelon abroad. However, several watermelon diseases reduce the yield and quality of this fruits and hampering this potential export market. Therefore, the main aims of this study are to identify the diseases of watermelons with their causal organisms and to evaluate their prevalence.

## Materials and methods

### Study site

A case study on watermelon diseases was conducted across eight blocks of four Upazilas in Sylhet and Natore Districts of Bangladesh in 2024. These two districts were considered due to their contrasting distinct characteristics in climatic conditions and soil properties [11–12]. Sylhet has the lowest temperature and highest rainfall while Natore has the lowest rainfall and high temperature (Table 1). However, molecular analysis was performed in the Plant Pathology Lab at Sher-e-Bangla Agricultural University in Dhaka.

**Table 1:**
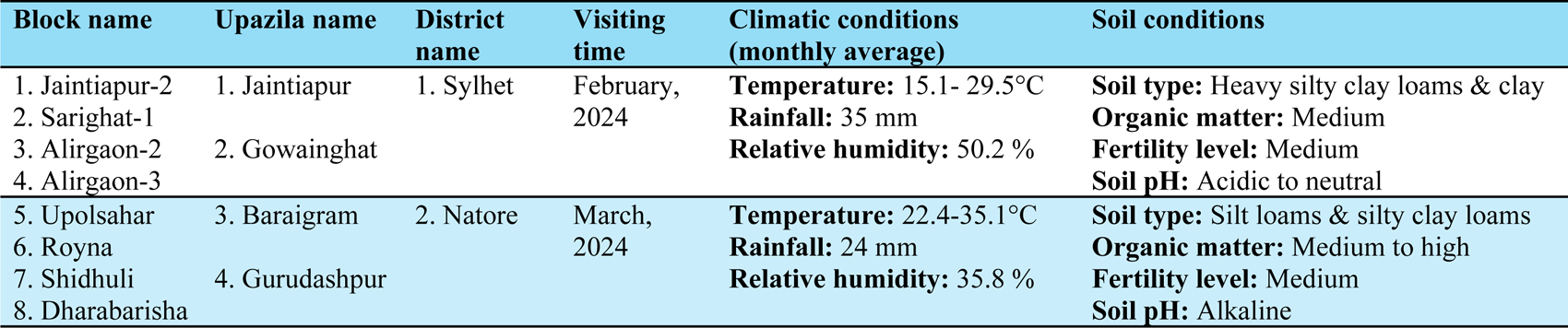
Field survey locations and environmental conditions for watermelon disease assessment in Bangladesh, 2024.

### Data collection

In total 40 fields from 8 blocks, (5 fields from each block) were examined and data were recorded on disease symptoms, affected plant parts, intensity, status, and disease occurrence percentage. Visiting each watermelon field, 10 plants were selected (total 400 plants from 40 fields) randomly to evaluate the disease occurrence. Disease occurrence was calculated employing the subsequent formula [13]:

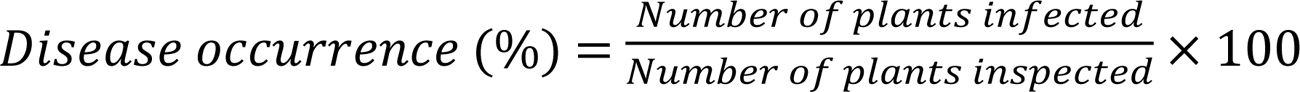

### Sample collection

Fungal and bacterial disease-infected plant parts (leaves, fruit, twigs, stem) exhibiting obvious symptoms were collected from watermelon disease surveyed fields. The collected specimens were stored in sterile zipper bags within an insulated ice box, labeled, and shifted to the Plant Pathology laboratory.

### Lab experiment

In the laboratory, the samples were stored at 4°C. The tissue planting method was employed, involving careful slicing of plant sections into small fragments, followed by rinsing and surface sterilization with 1% Mercuric Chloride (HgCl2). The sterilized fragments were then placed on Potato Dextrose Agar (PDA) medium and incubated for 6-7 days at 25±2°C. After that, the fungal and bacterial colonies were transferred to new PDA and NA (Nutrient Agar) media respectively, to establish pure cultures. The obtained pure cultures were stored at -20°C in a refrigerator for further analysis.

### Molecular analysis

The molecular detection process at the Plant Pathology Laboratory comprised DNA extraction, quantification, PCR amplification, gel electrophoresis with documentation, DNA purification, sequencing, and bioinformatics analysis.

### Extraction of genomic DNA

**For fungi:** DNA was extracted from plant tissues directly, infected by fungi and obligate pathogen viz. *Pseudoperonospora cubensis* (cannot be cultured on artificial media). In this step, the sample with a large lesion, and several conidia, mycelia, or sporangia were selected to extract DNA following the cetyltrimethylammonium bromide (CTAB) method as described in [14].

**For bacteria:** In the case of bacteria, overnight cultures on NA media were performed in the conventional and widely used method of DNA isolation based on phenol-chloroform extraction using centrifugation at 15,000 × g (Model: Z-216 M, HERMLE, Germany) as performed by [15–16]. The obtained DNA pellets were then stored at -20 °C before further analysis.

### DNA quantification

The NanoDrop Spectrophotometer (Model: ND2000, Thermo Scientific, USA) assessed DNA purity and concentration by measuring absorbance at 260/280 nm [17].

### Polymerase chain reaction

For fungus, PCR amplification of the ITS1 region utilized primers ITS1 (5’-TCCGTAGGTGAACCTGCGG-3’) and ITS4 (TCCTCCGCTTATTGATATGC-3’) [18].

Reactions were conducted in a Biorad thermocycler (S 1000TM), with an initial denaturation step of 5 minutes at 94°C, followed by 40 cycles of denaturation at 94°C for 1 minute, primer annealing at 53°C for 45 seconds, and primer extension at 72°C for 90 seconds. The reaction mix contained 25 µL, including 2 µL (20 ng/ml) DNA templates, 12.5 µL master mix, and 10 pmol of each primer.

In the case of bacteria, the 16S ribosomal RNA (rRNA) gene was amplified using the bacteria-specific primer 27F (5′-AGAGTTGATCCTGGCTCAG-3′) and universal primer 1492R (5′-GGTTACCTTGTTACGACTT-3′), generating 1465-bp amplicons [19]. The PCR reaction mixture, totaling 25 μL, included 12.5 μL of GoTaq™ G2 hot start master mix, 1 μL of DNA (25-65 ng/μl), 1 μL of Primer 27F (10-20 pMol), 1 μL of Primer 1492R (10-20 pMol), and 9.5 μL of nuclease-free water [20]. PCR was conducted in a Gene Atlas automated thermal cycler (Model: G2) with an initial denaturation at 95 °C for 5 min, followed by 30 cycles at 95 °C for 30 s, 52 °C for 45 s, and 72 °C for 90 s, with a final extension at 72 °C for 10 min, and an overnight hold at 4°C [21]. A 5 μL aliquot of each PCR product was electrophoresed on a 1% agarose gel and stored at -20°C.

### Electrophoresis and gel documentation

Agarose powder (Cat: V3125) was dissolved in Tris Borate EDTA (TBE) buffer (Cat: V4251) at 80°C for 5 min. Ethidium bromide (Cat: H5041) was added for DNA binding [21]. The gel was cast onto a Horizontal Gel Electrophoresis apparatus (Model: Mini, CBS Scientific, USA). DNA samples and a 1 kb ladder (Cat: G754B) were loaded into wells. Electrophoresis at 90 volts for 30 min separated DNA fragments by size. Gel documentation was performed using an Alpha Imager Gel Documentation System (Model: Mini, Protein Simple, USA) for visualization and image capture.

### DNA purification

DNA bands from the gel were dissolved in Membrane Binding Solution, mixed with PCR amplification products, and transferred to SV Minicolumns for binding. Subsequent washing with Membrane Wash Solution containing ethanol removed impurities. After ethanol evaporation, elution was performed by transferring the Minicolumns to clean tubes, adding nuclease-free water, and centrifuging to obtain purified DNA for further applications.

### DNA sequencing

Purified DNA samples were sent to Bioneer (Seoul, Korea) for partial sequencing via Sanger sequencing analysis. The resulting Sanger sequences were processed using Chromas 2.6. software to generate a FASTA file containing the partial sequence.

### Sequence analysis

FASTA files of all the obtained sequences were analyzed through nucleotide BLAST by submitting the sequences in the NCBI database (https://blast.ncbi.nlm.nih.gov/Blast.cgi) to match with existing sequences in the GenBank and obtained accession numbers. After that, the evolutionary history was inferred by using the Maximum Likelihood method and the Tamura-Nei model [22]. The bootstrap consensus tree inferred from 1000 replicates was taken to represent the evolutionary history of the taxa analyzed [23]. Branches corresponding to partitions reproduced in less than 50% of bootstrap replicates were collapsed. Initial tree(s) for the heuristic search were obtained automatically by applying Neighbor-Join and BioNJ algorithms to a matrix of pairwise distances estimated using the Tamura-Nei model and then selecting the topology with superior log likelihood value. Codon positions included were 1st+2nd+3rd+Noncoding. These analyses involved 18 nucleotide sequences of fungal and water mold and 9 sequences of bacteria. Evolutionary analyses were performed in MEGA11 software [24].

## Results

### Identification of diseases

During the survey, ten types of watermelon diseases were observed visually and initially confirmed by their distinctive characteristic symptoms described in (Table 2). Among the identified diseases were four fungal, three bacterial, two water mold, and one viral disease. All of the symptoms were observed above the ground parts of the watermelon plant especially leaves, fruits, stems, etc. shown in (Figure 1).

**Figure 1.**
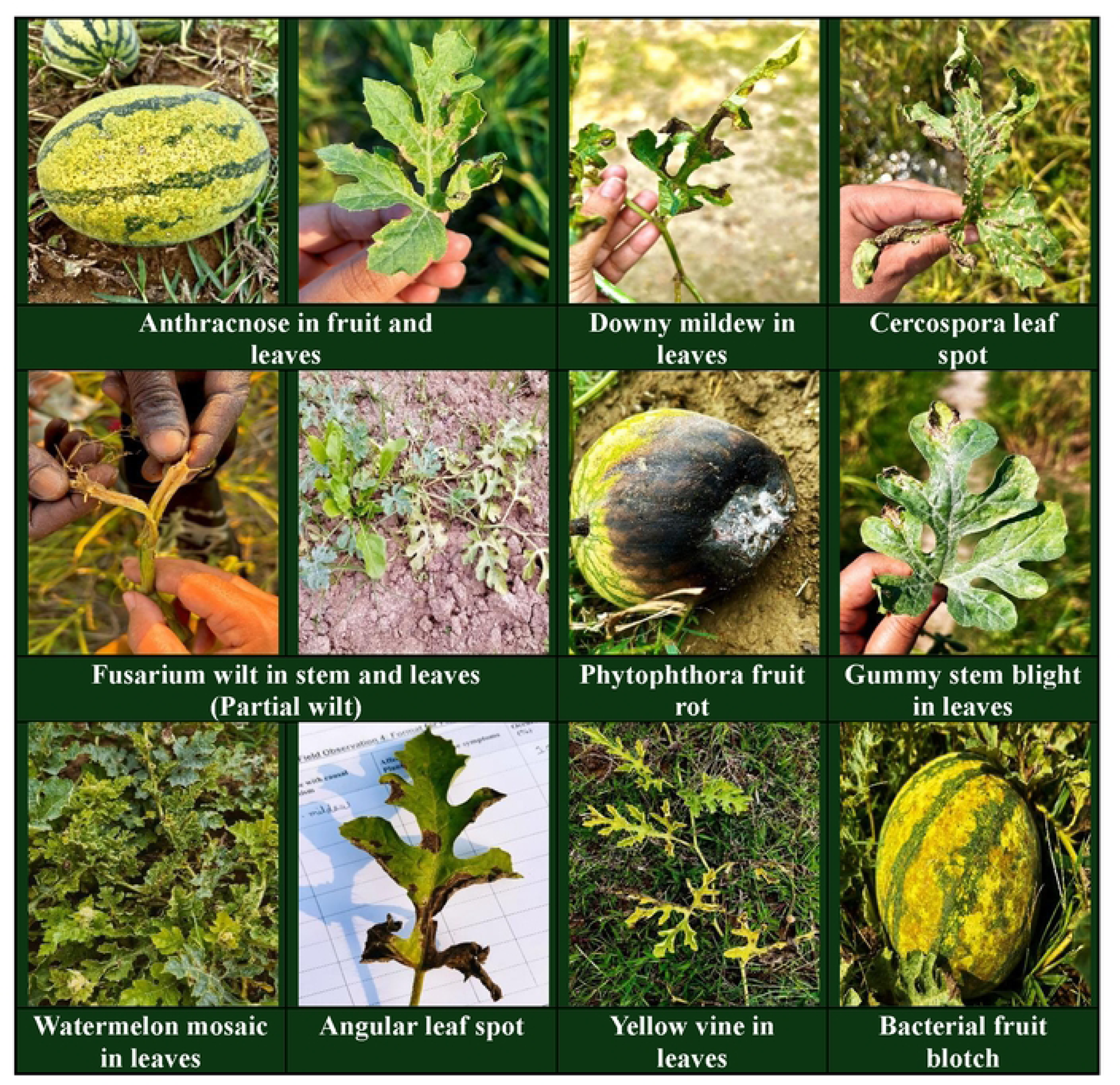
Watermelon disease symptoms were observed in different infected plant parts during the field study.

**Table 2.**
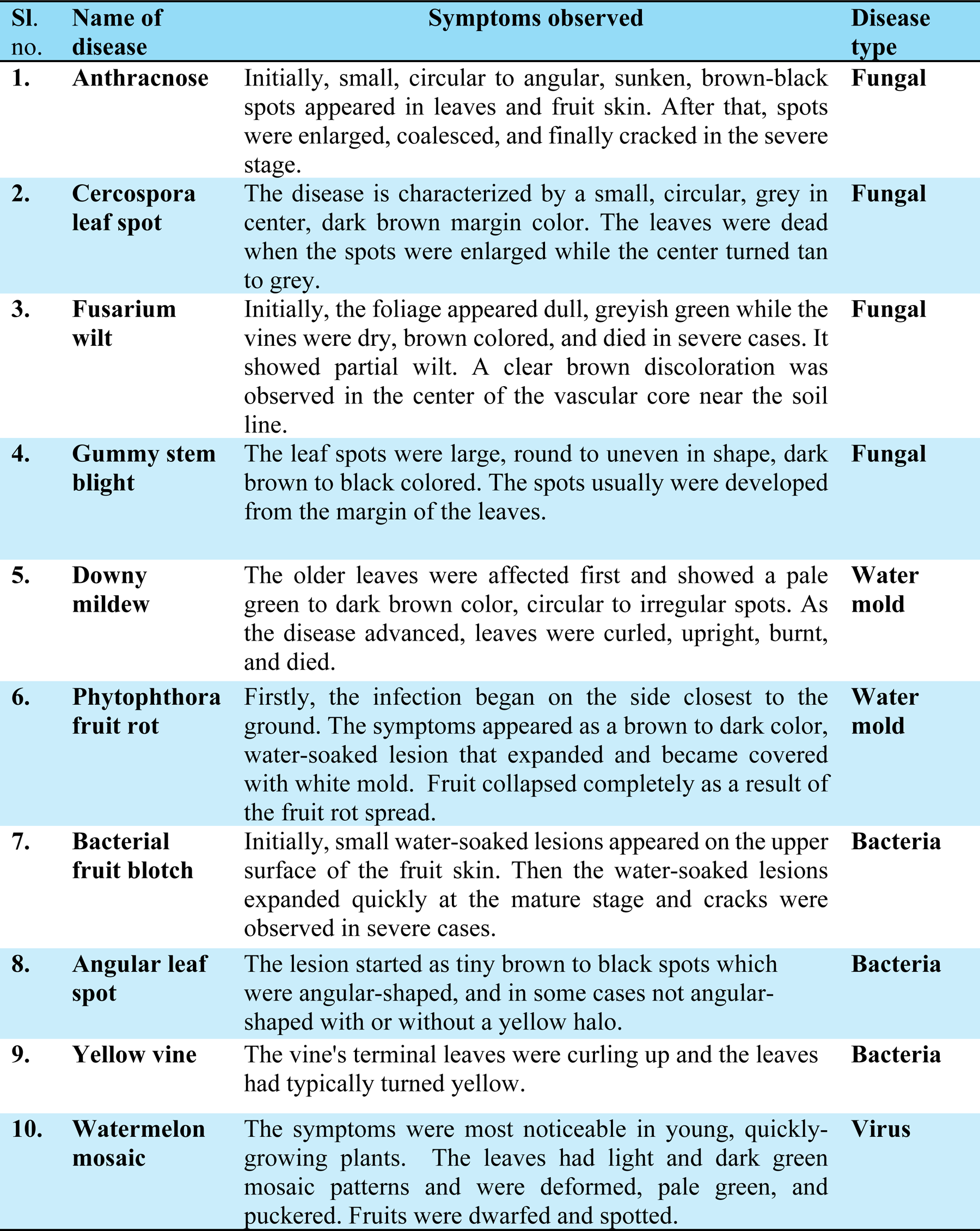
Symptoms of watermelon diseases were observed under field conditions in the Sylhet and Natore districts of Bangladesh.

### Identification of the causal organisms

A molecular approach was performed to identify the causal organisms of watermelon diseases in the lab where nine species of the pathogens were identified except the viral pathogen. The identified pathogens were *Colletrotrichum orbiculare* isolate NatBD-6 (Anthracnose), *Cercospora citrullina* isolate NatBD-7 (Cercospora leaf spot), *Fusarium oxysporum* isolate isolate NatBD-8 (Fusarium wilt), *Stagonosporopsis cucurbitacearum* isolate SylBD-1 (Gummy stem blight), *Pseudoperonospora cubensis* isolate SylBD-2 (Downy mildew), *Phytophthora capsici* isolate SylBD-3 (Phytophthora fruit rot), *Acidovorax citrulli* isolate SylBD-4 (Bacterial fruit blotch), *Pseudomonas syringae* isolate SylBD-5 (Angular leaf spot), and *Serratia marcescens* isolate NatBD-9 (Yellow vine).

### Study on PCR products

The Gel-Doc system visualized PCR products of fungal and water mold isolates generated by ITS1 and ITS4 primers, resulting in the DNA bands of around 710 base pairs (bp). Variability in amplicon sizes ranging from 512 to 751 bp was observed in the GenBank database. In the case of bacterial isolates, the DNA bands of approximately 1465 bp were detected using primers 27F and 1492R where the amplicon sizes were 1367 to 1410 bp in the database. The generated DNA bands indicated the successful PCR amplification, which were extremely similar in size to the target band. The size of the amplified DNA fragment was verified using a Bench Top 1 kb DNA ladder, which acts as a size marker. Figure 2 (Lanes 1-6) shows the successful amplification of DNA bands obtained from several fungal and water mold isolates whereas Figure 3 (Lanes 1-3), displays the bacterial isolates. These confirm the specificity and successful amplification of the desired DNA fragments under the utilized primer and PCR conditions, as demonstrated by the Gel-Doc system.

**Figure 2.**
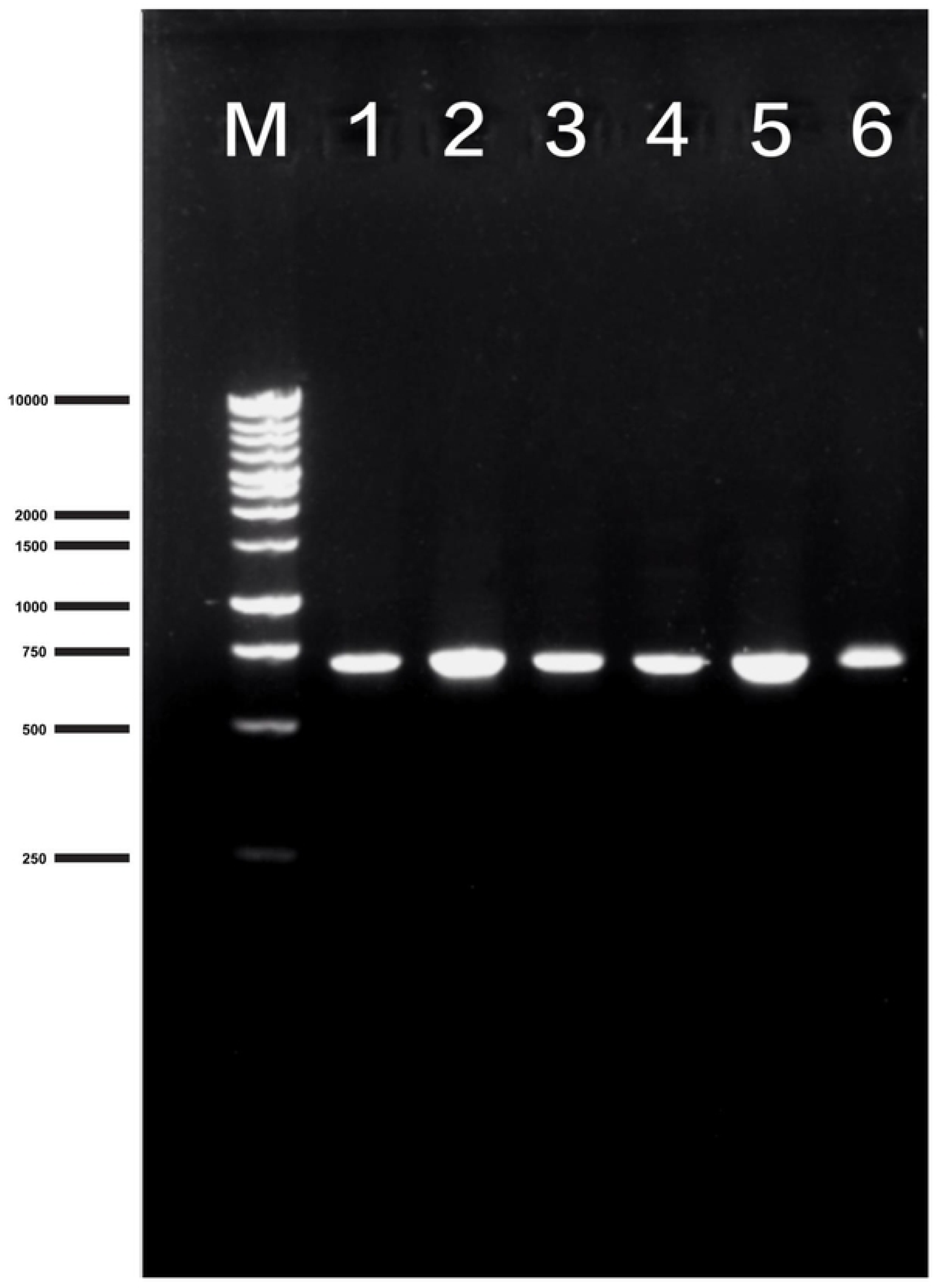
PCR amplified products of fungus and water mold obtained from (1) *Colletrotrichum orbiculare* isolate NatBD-6, (2) *Cercospora citrullina* isolate NatBD-7, (3) *Fusarium oxysporum* isolate NatBD-8, (4) *Stagonosporopsis cucurbitacearum* isolate SylBD-1, (5) *Pseudoperonospora cubensis* isolate SylBD-2, and (6) *Phytophthora capsici* isolate SylBD-3. M: denotes 1kb DNA ladder (Marker).

**Figure 3.**
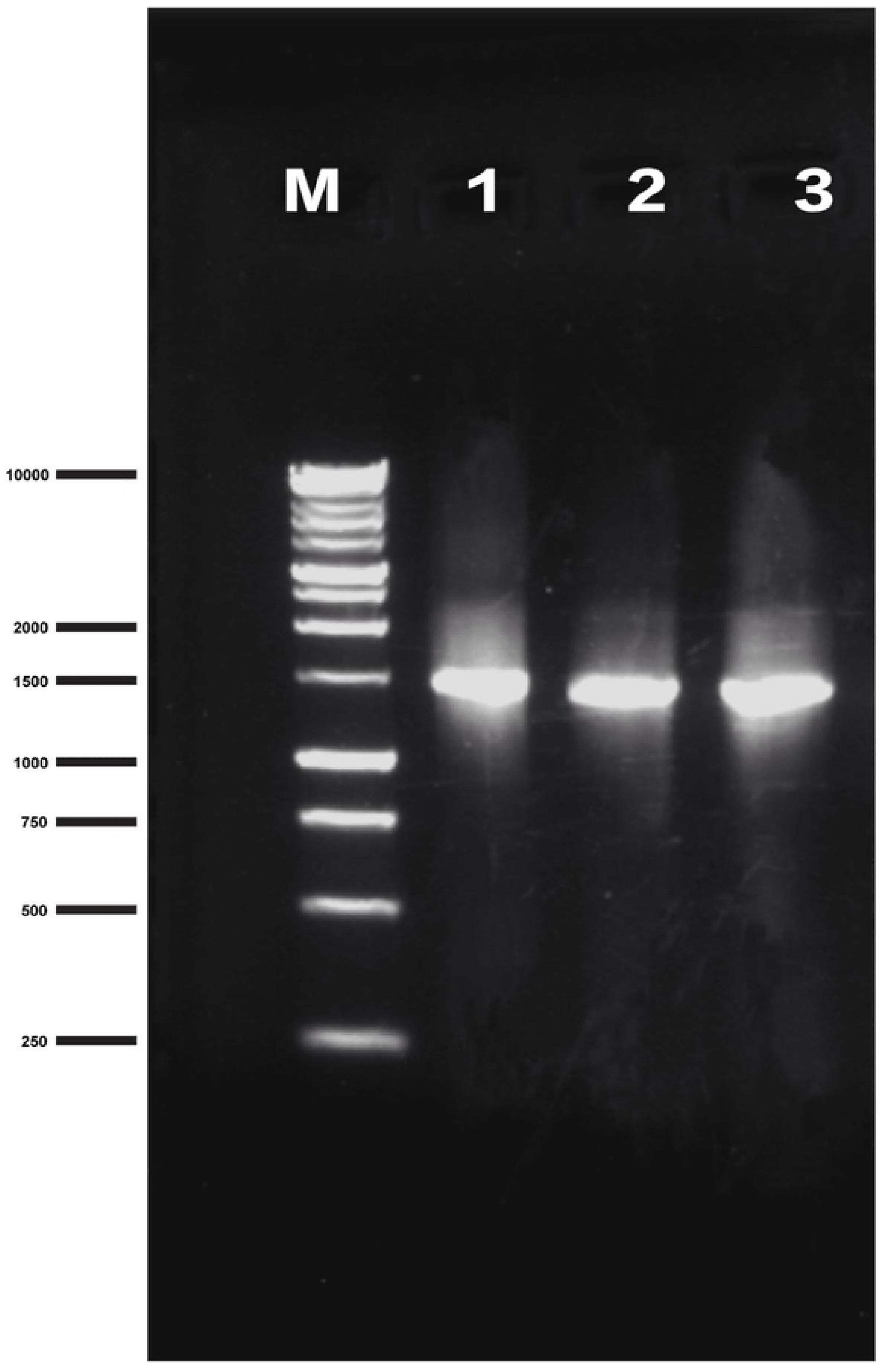
PCR amplified products of bacteria obtained from (1) *Acidovorax citrulli* isolate SylBD-4, (2) *Pseudomonas syringae* isolate SylBD-5, and (3) *Serratia marcescens* isolate NatBD-9. M: denotes 1kb DNA ladder (Marker).

### Analysis of DNA sequences

The obtained FASTA sequences underwent analysis using the BLAST tool on the NCBI website (S1 Appendix). They were revealed matches with existing nucleotide sequences in the NCBI GenBank database. All the sequences exceeding 98% (98.91-99.71%) homology with their corresponding sequence. Subsequently, unique accession numbers were assigned to the partial sequences for four fungal, two water mold, and three bacterial isolates (Table 3).

**Table 3:** Results of gene identification with the BLAST Program.

| LAB strain |  |  | BLAST Alignment |  |  |  |  |
| --- | --- | --- | --- | --- | --- | --- | --- |
| Sl. no. | Identified pathogen isolates | Obtained accession no. | Species | Query coverage | E value | Percent identity | Accession no. |
| 1. | *Colletotrichum orbiculare* isolate NatBD-6 | PP837831 | *Colletotrichum orbiculare* isolate POL1 | 100% | 0.0 | 99.26% | ON398802 |
| 2. | *Cercospora citrullina* isolate NatBD-7 | PP837832 | *Cercospora citrulline* isolate Cer 76-18 | 100% | 0.0 | 98.91% | OL589543 |
| 3. | *Fusarium oxysporum* isolate NatBD-8 | PP837833 | *Fusarium oxysporum* f. sp. *niveum* | 100% | 0.0 | 99.39% | FJ156282 |
| 4. | *Stagonosporopsis cucurbitacearum* isolate SylBD-1 | PP837834 | *Stagonosporopsis cucurbitacearum* isolate RKS8 | 100% | 0.0 | 99.22% | OL774657 |
| 5. | *Pseudoperonospora cubensis* isolate SylBD-2 | PP837835 | *Pseudoperonospora cubensis* isolate E1CHK0 | 100% | 0.0 | 99.18% | MW845754 |
| 6. | *Phytophthora capsici* isolate SylBD-3 | PP837836 | *Phytophthora capsici* isolate 219 | 100% | 0.0 | 99.60% | DQ464028 |
| 7. | *Acidovorax citrulli* isolate SylBD-4 | PP837828 | *Acidovorax citrulli* strain RKFB 793 | 100% | 0.0 | 99.71% | KP410333 |
| 8. | *Pseudomonas syringae* isolate SylBD-5 | PP837829 | *Pseudomonas syringae* strain BP550 | 100% | 0.0 | 99.57% | KX533931 |
| 9. | *Serratia marcescens* isolate NatBD-9 | PP837830 | *Serratia marcescens* strain QQ3 | 100% | 0.0 | 99.57% | OP861103 |

### Analysis of phylogenetic tree

The phylogenetic trees were analyzed using the sequences of fungal, water mold, and bacterial sequences where the highest log likelihood was -4119.79 for fungal and water mold sequences. At the same time, for bacterial sequences, it was log -3797.87. There were a total of 1065 and 1446 positions in the final dataset obtained from fungal and bacterial sequences respectively. Based on the phylogenetic tree, the studied strains fall into six main clusters of fungal and water mold and three main clusters of bacterial isolates. In (Figure 4), shows the clusters of *Colletrotrichum, Fusarium, Pseudoperonospora, Cercospora, Stagonosporopsis, and Phytophthora* where all the clusters exhibited 100% of replicate trees in which the associated taxa clustered together in the bootstrap test 1000 replicates are shown next to the branches. On the other hand, (Figure 5) shows the clusters of bacterial isolates which were *Acidovorax, Pseudomonas,* and *Serratia* and these displayed 100% replicate trees for all the clusters. The genetic distances between the strains are also shown in the trees. The distances are measured by the number of substitutions per site. Considering the percentage of replicate trees and genetic distances, the phylogenetic tree shows that the studied strains were all relatively closely related to each other within a cluster.

**Figure 4.**
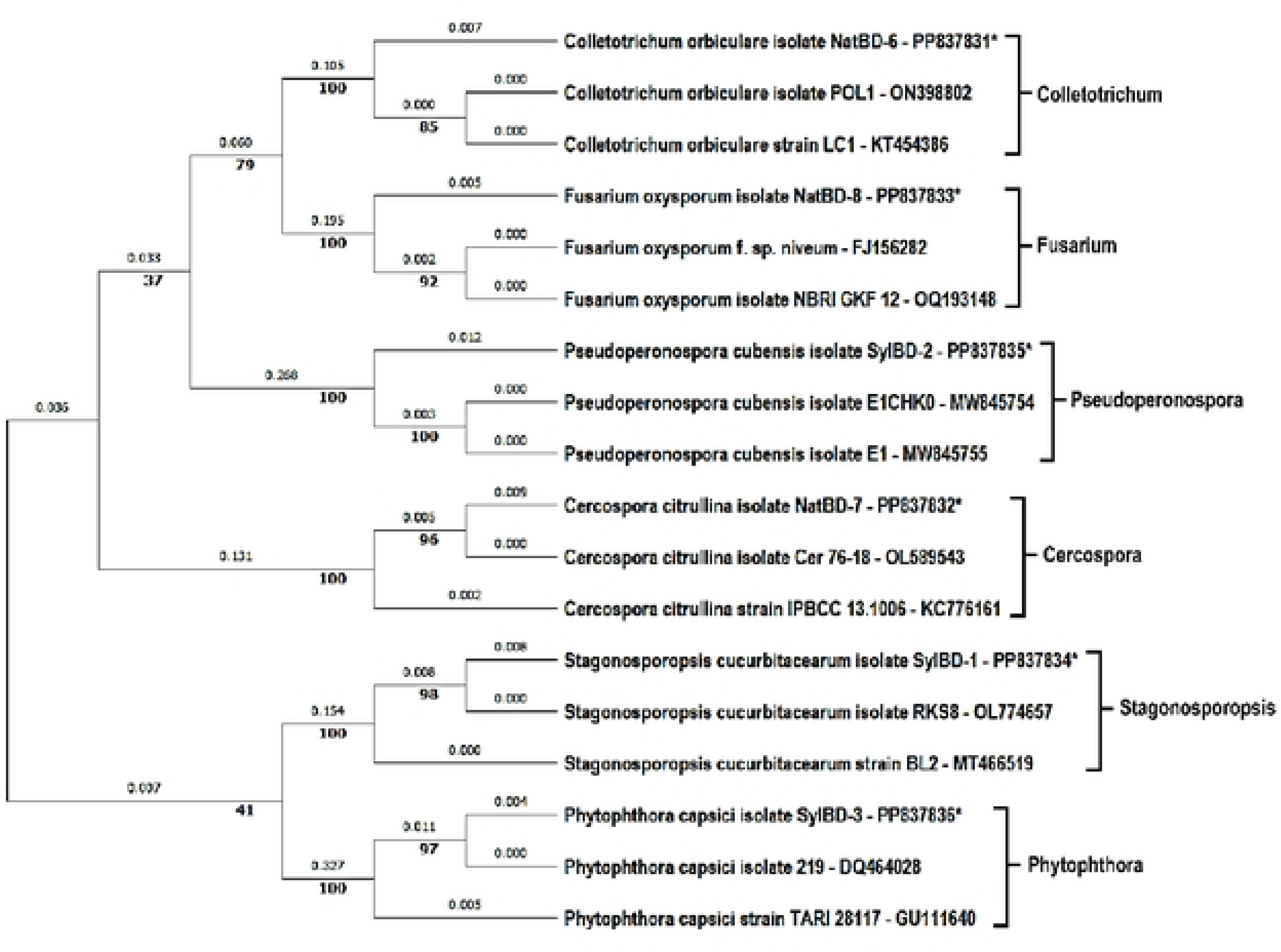
Phylogenetic tree for fungal and water mold pathogens of watermelon plants collected from Natore and Sylhet regions in Bangladesh. This analysis involved 18 nucleotide sequences where the clusters fall into 6 distinct clades of fungal and water mold strains. * Denotes the isolates obtained from the watermelon samples.

**Figure 5.**
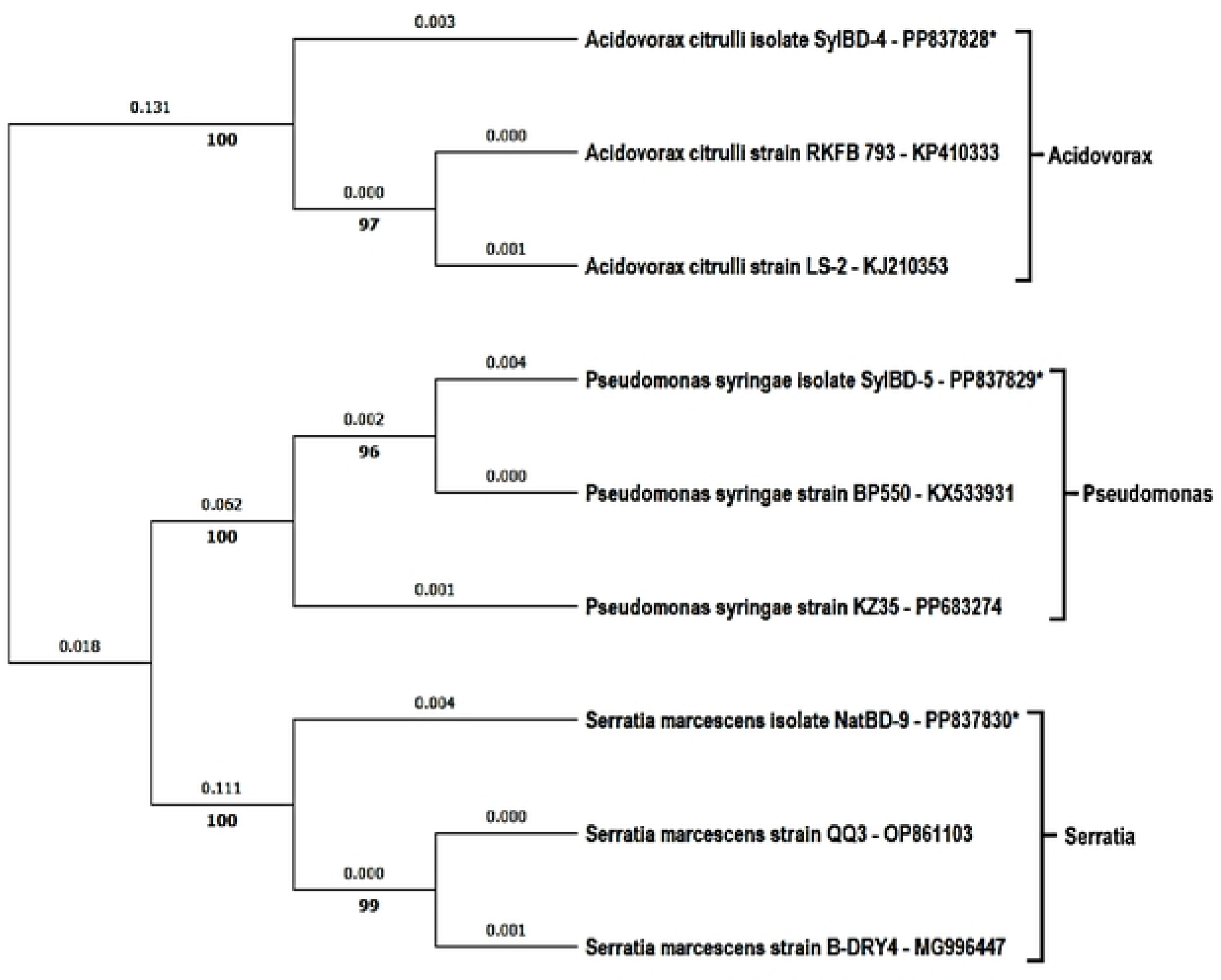
Phylogenetic tree for bacterial pathogens of watermelon plants collected from Natore and Sylhet regions in Bangladesh. This analysis involved 9 nucleotide sequences where the clusters fall into 3 distinct clades of bacterial strains. * Denotes the isolates obtained from the watermelon samples.

### Evaluation of disease status, intensity, and occurrence

During the survey, 400 plants were observed in 40 fields of 8 different blocks in Sylhet and Natore districts to evaluate the prevalence of corresponding diseases (S2 Appendix). Among the identified diseases, the highest disease occurrence in Sylhet was Fusarium wilt (47.58%) followed by Gummy stem blight (41.5), and in Natore was Angular leaf spot (37.5) followed by Yellow vine (33%). However, Downy mildew and Watermelon mosaic diseases were completely absent in the Natore region while Cercospora leaf spot disease was not observed in the Sylhet zone. In the early growing stage of watermelon plants, Fusarium wilt showed the most intent to damage the plants and spread as a major disease while Gummy stem attacked in the late stage with medium disease intensity. However, Phytophthora fruit rot was evaluated as a minor disease but its intensity to damage the watermelon fruits was high (Table 4).

**Table 4.**
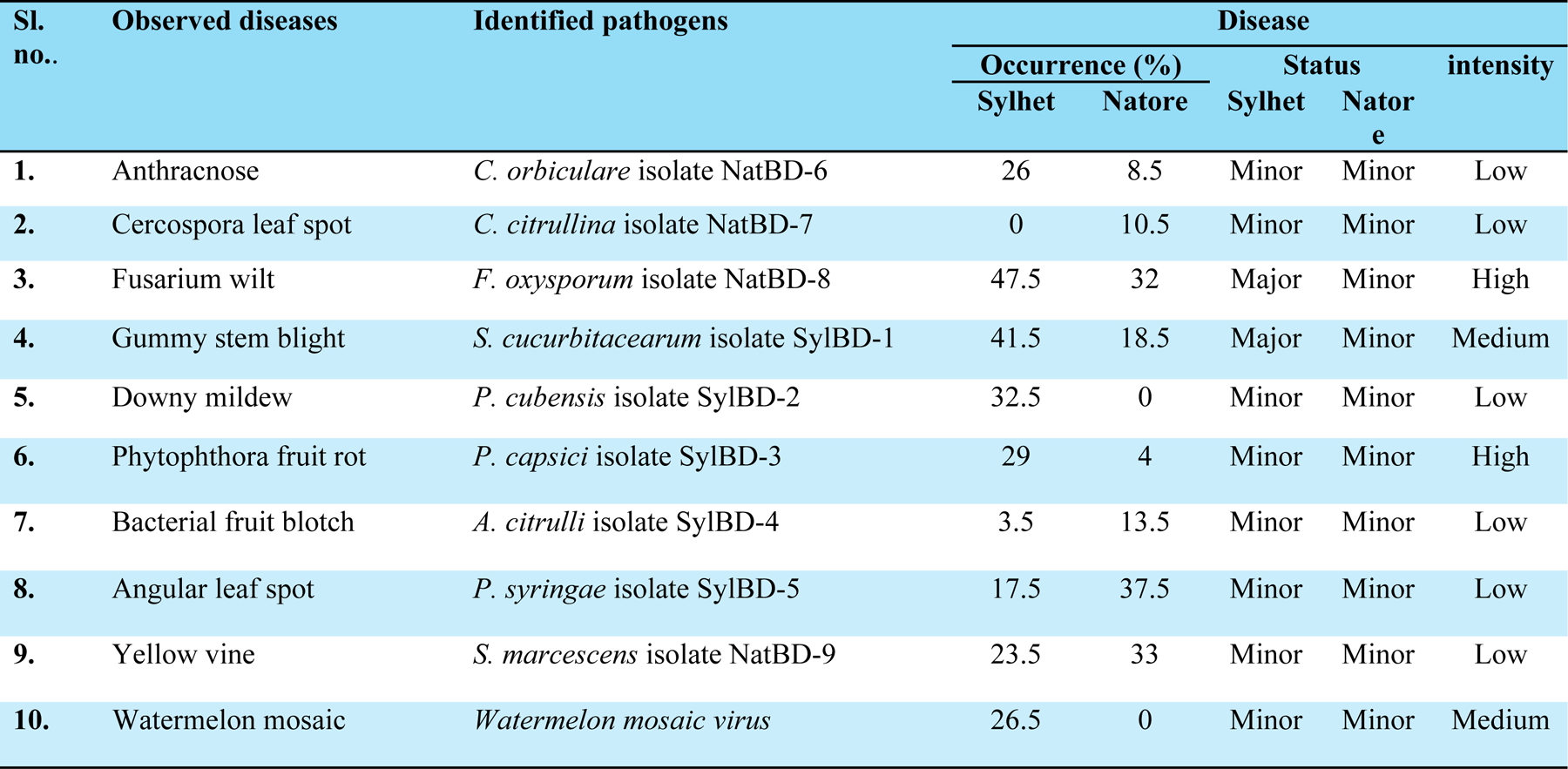
Prevalence of the identified watermelon diseases in Sylhet and Natore regions.

| Sl. no.. | Observed diseases | Identified pathogens | Disease |  |  |  |  |
| --- | --- | --- | --- | --- | --- | --- | --- |
|  |  |  | Occurrence (%) |  | Status |  | intensity |
|  |  |  | Sylhet | Natore | Sylhet | Natore |  |
| 1. | Anthracnose | *C. orbiculare* isolate NatBD-6 | 26 | 8.5 | Minor | Minor | Low |
| 2. | Cercospora leaf spot | *C. citrullina* isolate NatBD-7 | 0 | 10.5 | Minor | Minor | Low |
| 3. | Fusarium wilt | *F. oxysporum* isolate NatBD-8 | 47.5 | 32 | Major | Minor | High |
| 4. | Gummy stem blight | *S. cucurbitacearum* isolate SylBD-1 | 41.5 | 18.5 | Major | Minor | Medium |
| 5. | Downy mildew | *P. cubensis* isolate SylBD-2 | 32.5 | 0 | Minor | Minor | Low |
| 6. | Phytophthora fruit rot | *P. capsici* isolate SylBD-3 | 29 | 4 | Minor | Minor | High |
| 7. | Bacterial fruit blotch | *A. citrulli* isolate SylBD-4 | 3.5 | 13.5 | Minor | Minor | Low |
| 8. | Angular leaf spot | *P. syringae* isolate SylBD-5 | 17.5 | 37.5 | Minor | Minor | Low |
| 9. | Yellow vine | *S. marcescens* isolate NatBD-9 | 23.5 | 33 | Minor | Minor | Low |
| 10. | Watermelon mosaic | *Watermelon mosaic virus* | 26.5 | 0 | Minor | Minor | Medium |

## Discussion

This study aimed to investigate watermelon diseases across two distinct agro-ecological zones in Bangladesh, Sylhet, and Natore, providing valuable insights into these diseases’ prevalence, causative agents, and molecular identification. The contrasting climatic conditions of these districts Sylhet with its lower temperatures and high rainfall, and Natore with higher temperatures and low rainfall served as a compelling backdrop for understanding disease dynamics in varied environmental contexts.

### Disease Prevalence

The survey identified ten diseases affecting watermelon plants, categorized into fungal, bacterial, water mold, and viral diseases. Notably, Fusarium wilt emerged as the most prevalent disease, with an occurrence rate of 47.5%. This high prevalence aligns with global reports highlighting Fusarium wilt as a significant threat to watermelon production due to its soil-borne nature and persistent survival in soil [25]. Gummy stem blight and Angular leaf spot were also prevalent, with occurrences of 41.5% and 37.5%, respectively [26–27]. The survey results underscore the need for targeted management strategies, particularly for Fusarium wilt, which exhibited the highest intensity during the early growth stages of watermelon plants.

### Identification of Pathogens

The molecular identification, including DNA extraction, PCR amplification, gel electrophoresis, and sequencing was crucial in accurately identifying the pathogens. The study’s use of specific primers for fungal (ITS1/ITS4) and bacterial (27F/1492R) DNA regions ensured precise amplification of target sequences. The molecular analysis process confirmed nine distinct pathogen species responsible for the observed diseases where the sequencing revealed high homology (98.91-99.71%) with known sequences in the GenBank database, affirming the accuracy of pathogen identification. The pathogens identified included *Colletrotrichum orbiculare* NatBD-6 (Anthracnose), *Cercospora citrullina* NatBD-7 (Cercospora leaf spot), *Fusarium oxysporum* NatBD-8 (Fusarium wilt), *Stagonosporopsis cucurbitacearum* SylBD-1 (Gummy stem blight), *Pseudoperonospora cubensis* SylBD-2 (Downy mildew), *Phytophthora capsici* SylBD-3 (Phytophthora fruit rot), *Acidovorax citrulli* SylBD-4 (Bacterial fruit blotch), *Pseudomonas syringae* SylBD-5 (Angular leaf spot), and *Serratia marcescens* NatBD-9 (Yellow vine). The identification of these pathogens also identified by several researchers in the watermelon field provides a crucial foundation for developing specific control measures [5–6].

### Phylogenetic Analysis

The phylogenetic analysis revealed six main clusters for fungal and water mold pathogens and three for bacterial isolates, indicating the genetic relatedness among the strains. The high bootstrap values (100%) in the phylogenetic trees confirm the reliability of these groupings, providing a robust framework for understanding the evolutionary relationships of the pathogens involved.

### Environmental Influence on Disease Distribution

The contrasting environmental conditions of Sylhet and Natore significantly influenced disease distribution and prevalence. The high rainfall and cooler temperatures in Sylhet likely favored the proliferation of water molds and certain fungal pathogens, such as *Pseudoperonospora cubensis* and *Phytophthora capsica, Colletotrichum orbiculare*. Conversely, the hotter and drier conditions in Natore were more conducive to bacteria like *Acidovorax citrulli* and *Pseudomonas syringae*.

This environmental influence underscores the necessity of region-specific disease management practices that consider local climatic conditions.

### Implications for Disease Management

The findings of this study have important implications for watermelon disease management in Bangladesh. The high prevalence and intensity of Fusarium wilt, particularly in the early growth stages, highlight the need for early intervention strategies, such as soil fumigation, crop rotation, and resistant cultivars. For other prevalent diseases like Gummy stem blight and Angular leaf spot, integrated pest management (IPM) approaches, including the use of fungicides, proper irrigation practices, and resistant varieties, should be emphasized. The molecular identification of pathogens also opens avenues for the development of molecular-based diagnostic tools, enabling rapid and accurate detection of these pathogens in the field.

## Conclusion

This comprehensive study provides a detailed account of the prevalence, identification, and molecular characterization of watermelon diseases in two agroecologically distinct districts of Bangladesh. The findings highlight the significant impact of environmental conditions on disease distribution and underscore the need for tailored disease management strategies. Future research should focus on developing and implementing integrated disease management practices that consider local environmental conditions and pathogen characteristics. Such efforts will be crucial in mitigating the impact of these diseases on watermelon production and ensuring sustainable agricultural practices in the region.

## Compliance with ethical standards

### Ethical approval

A total of 40 fields were permitted for collection of plant-infected parts samples. Department of Agricultural Extension, Govt. of Bangladesh granted the permission to perform this project. Sample collection was done with the permission and presence of all the corresponding farmers in their fields.

### Data Availability Statement

All relevant data are within the manuscript and its Supporting Information files.

## Acknowledgments

The author would like to thank the Center for Resource Development Studies Ltd. (CRDS) for allowing him to do the field survey in the Sylhet and Natore regions.

## Funding

The fund was allocated by the Center for Resource Development Studies Ltd. only for field investigation, no external fund was received.

### Author contribution statement

RF = Conceptualization, Data curation, Formal analysis, Funding acquisition, Investigation, Methodology, Project administration, Resources, Software, Validation, Visualization, Writing - original draft, review, and editing.

